# Differential virulence of *Trypanosoma brucei rhodesiense* isolates does not influence the outcome of treatment with anti-trypanosomal drugs in the mouse model

**DOI:** 10.1101/2020.01.30.926675

**Authors:** Kariuki Ndung’u, Grace Adira Murilla, John Kibuthu Thuita, Geoffrey Njuguna Ngae, Joanna Eseri Auma, Purity Kaari Gitonga, Daniel Kahiga Thungu, Richard Kiptum Kurgat, Judith Kusimba Chemuliti, Raymond Ellie Mdachi

## Abstract

We assessed the virulence and anti-trypanosomal drug sensitivity patterns of *Trypanosoma brucei rhodesiense* (*Tbr*) isolates in the Kenya Agricultural and Livestock Research Organization-Biotechnology Research Institute (KALRO-BioRI) cryobank. Specifically, the study focused on *Tbr* clones originally isolated from the western Kenya/eastern Uganda focus of human African Trypanosomiasis (HAT). Twelve (12) *Tbr* clones were assessed for virulence using groups(n=10) of Swiss White Mice monitored for 60 days post infection (dpi). Based on survival time, four classes of virulence were identified: (a) very-acute: 0-15, (b) acute: 16-30, (c) sub-acute: 31-45 and (d) chronic: 46-60 dpi. Other virulence biomarkers identified included: prepatent period (pp), parasitaemia progression, packed cell volume (PCV) and body weight changes. The test *Tbr* clones together with KALRO-BioRi reference drug-resistant and drug sensitive isolates were then tested for sensitivity to melarsoprol (mel B) pentamidine, diminazene aceturate and suramin, using mice groups (n= 5) treated with single doses of each drug at 24 hours post infection. Our results showed that the clones were distributed among four classes of virulence as follows: 3/12 (very-acute), 3/12 (acute), 2/12 (sub-acute) and 4/12 (chronic) isolates. Differences in survivorship, parasitaemia progression and PCV were significant (P<0.001) and correlated. The isolate considered to be drug resistant at KALRO-BioRI, KETRI 2538, was confirmed to be resistant to melarsoprol, pentamidine and diminazene aceturate but it was not resistant to suramin. At least 80% cure rates of all the test isolates was achieved with melarsoprol (1mg/Kg and 20 mg/kg), pentamidine (5 and 20 mg/kg), diminazene aceturate (5 mg/kg) and suramin (5 mg/kg) indicating that the isolates were not resistant to any of the drugs despite the differences in virulence. This study provides evidence of variations in virulence of *Tbr* isolates from a single HAT focus and confirms that these variations are not a significant determinant of isolate sensitivity to anti-trypanosomal drugs.

## Introduction

Human African trypanosomiasis (HAT), also known as sleeping sickness, is a vector-borne parasitic disease. It is caused by infection of humans with protozoan parasites belonging to the genus *Trypanosoma*. HAT is caused by two species of trypanosomes, namely *Trypanosoma brucei gambiense* and *Trypanosoma brucei rhodesiense* [1]. They are transmitted to humans by tsetse fly (*Glossina* genus)[2]. *Trypanosoma brucei gambiense* is found in countries in West and Central Africa and causes a chronic infection [3]. A person can be infected for months or even years without major signs or symptoms of the disease [4]. When more evident symptoms emerge, the patient is often already in an advanced disease stage where the central nervous system is affected. *Trypanosoma brucei rhodesiense* is found in countries in eastern and southern Africa and causes an acute infection Symptoms manifest within 2-4 weeks of infective bite [3].

HAT develops in two stages namely, early (hemolymphatic) and late (meningo-encephalitic) stage. In the early stage of the disease, parasites proliferate in the blood and lymphatic system while in the late stage, parasites penetrate the blood brain barrier (BBB) and persist and proliferate in the central nervous system (CNS), causing an encephalitic reaction that leads to death if untreated or inadequately treated [5]. For first stage infections, there are no specific clinical signs and symptoms in both forms of the disease; fever, headache and loss of appetite are common [1] as well anemia in the monkey model [6]. With *T.b. rhodesiense infections*, first signs and symptoms are observed a few weeks after infection;[1]. However, a mild form of chronic *T. b. rhodesiense* infections with incubation times of several months has been reported in Zambia [7]. The acute and the chronic HAT infections caused by *T. b. rhodesiense* in different foci differs both in their inflammatory response and pathology. The pathology encountered in the acute HAT infections is characterized by elevated Tumor necrosis factor alpha (TNF-α) while that encountered in the chronic HAT infections is characterized by elevated transforming growth factor (TGF-β) [8].

Treatment of *Trypanosoma brucei rhodesiense* infections involves the use of early stage drugs such as pentamidine and suramin [9] and late stage drugs such as melarsoprol; melarsoprol is the only drug recommended by WHO for treatment of late-stage *T b rhodesiense* infection, but can be lethal to 5% of patients owing to post-treatment reactive encephalopathy [10]. HAT therapy is further complicated by reports of drug resistance in different foci, including against suramin and melarsoprol in Tanzania [11] and against melarsoprol in south Sudan [12]. In their study, [13] suggested that investigations into treatment failure in HAT and use of alternative drugs or treatment regimens should not only focus on differential genotypes of the parasites but also on differential virulence and tissue tropism as possible causes. The present study was therefore designed to investigate the occurrence of differential virulence of isolates recovered from Western Kenya/ Eastern Uganda HAT focus and the potential role of these variations on isolate sensitivity to anti-trypanosomal drugs. The study will also avail well characterized *T.b.rhodesiense* isolates for future studies.

## Materials and methods

### Ethics

All experimental protocols and procedures used in this study involving laboratory animals were reviewed and approved by Institutional Animal Care and Use Committee (IACUC) of Kenya Agricultural and Livestock Research Institute-Biotechnology Research Institute (KALRO-BioRI) Ref: C/Biori/4/325/II/53)

### Experimental animals

The study used 6–8 weeks old male Swiss White mice, each weighing 25–30 g live body weight. The animals were obtained from the Animal Breeding Unit at KALRO-BioRI, Muguga. The mice were housed in standard mouse cages and maintained on a diet consisting of commercial pellets (Unga^®^ Kenya Ltd). All experiments were performed according to the guidelines set by the Institutional Animal Care and Use Committee (IACUC) of KALRO-BioRI. Briefly, water was provided ad libitum. All mice were acclimatized for two weeks, during which time they were screened and treated for ecto and endoparasites using ivermectin (Ivermectin^®^, Anupco, Suffolk, England). During the two-week acclimation period, pre-infection data were collected on body weights and packed cell volume once a week prior to parasite inoculation.

### Trypanosomes and Cloning

Twelve *T.b. rhodesiense* trypanosome stabilates (KETRI 2482, 2487, 3304, 3305, 3380, 3664, 3798, 3800, 3801, 3803, 3926, 3928) were selected from the KALRO-BioRI specimen bank. Cloning was carried out as described by [14]. Briefly, the trypanosome stabilates were inoculated into mice immunosuppressed using cyclophosphamide at 100 mg/kg for three consecutive days (total dose 300mg/kg) body weight (bwt) as previously described [15]. Animals were monitored daily for parasitaemia. When the mice attained a parasitaemia score of 3.2.x10^7^ trypanosomes/mL [16], they were bled from the tail vein and the blood sample appropriately diluted using a mixture of PSG pH 8.0 and guinea pig serum in the ratio of 1:1. Using the hanging drop method [17], a single trypanosome was then picked using a syringe with a 25 gauge needle suspended in at least 0.2mls PSG pH 8.0 and injected intraperitoneally (ip) into a single immunosuppressed mouse. This was replicated ten times to increase the chances of success. Infected mice were then monitored for parasitaemia daily [16]. Any of the ten mice which became parasitaemic was euthanized using concentrated carbon dioxide, bled from the heart and the harvested trypanosomes cryopreserved in PSG pH 8.0 in 20% glycerol as a clone stabilate.

### Virulence studies

#### Design of virulence study

Male Swiss White mice were housed in groups of 10 in standard mouse cages containing wood shavings as bedding material. The cryopreserved cloned parasites were thawed, and injected ip into immunosuppressed donor Swiss White mice for multiplication. The mice were euthanized using carbon dioxide[18] at peak parasitaemia and blood collected from the heart in EDTA for quantification as previously described [19].The ten mice in each cage were infected with one *Tbr* clone, with each mouse receiving 1×10^4^ trypanosomes injected intraperitoneally. The infected mice were monitored for pre-patent period, parasitaemia progression, PCV, body weight and survival time as virulence biomarkers.

### Pre-patent period and Parasitaemia progression

Blood for estimation of parasitaemia levels was collected daily for the first 14 days and thereafter three times in a week from each mouse using the tail tip amputation method [20]. The PP and parasitaemia progression were determined using the rapid matching method of [16] [21]. The infected mice were monitored for 60 days post infection.

### Packed cell volume (PCV) and body weight changes

PCV was determined as outlined by [22]. Body weight (bwt) was measured once in a week using (Mettler Tolendo PB 302 ^®^, Switzerland) digital balance [19]

### Survival times and virulence classification

The classification of the trypanosome virulence were based on the survival of the infected mice as previously described [23]. The twelve *T.b. rhodesiense* clones were placed into four classes of virulence based on the survival of 60% or more of the infected mice as follows: very-acute (0-15 days), acute (16 – 30), sub-acute (31 – 45) and chronic classes (46 – 60). Each mouse’s survival time was based on attainment of the *at extremis* condition which was determined on a ≥ 25% drop in PCV and consistently high parasitaemia levels of 1×10^9^/ml for at least two consecutive days, [24]. The mice were euthanized by CO_2_ asphyxiation [25] and recorded as dead animal. Mice surviving at 60 dpi were equally euthanized, survival time recorded as 60 days and categorized as censored data.

### Drug sensitivity study

Initially, sensitivity patterns for KALRO-BiORI laboratory reference isolates considered drug resistant (KETRI 2538) or drug-sensitive (KETRI 3738) were determined Melarsoprol (Arsobal^®^, Aventis), Diminazene aceturate [(Veriben^®^,Ceva, France), Pentamidine (Pentacarinat^®^-Sanofi, UK) and Suramin (Germanin^^®^^ Bayer), using dose rates ranging from 1-40 mg/kg body weight (Table 2) in order to identify cut-off points for characterizing isolates as drug resistant. Thereafter, the *T. b. rhodesiense* test clones were evaluated for sensitivity to the same drugs (Table 3). An isolate was considered drug-resistant if 2/5 (40%) of the infected and treated mice relapsed [11] after having been treated at 20mg/kg bwt.

Suramin and Pentamidine drugs (100% w/w) for the highest dosage of 40mg/kg bw was prepared by dissolving 40mg of these drugs in 10mls distilled water to give a concentration of 4 mg/ml. Diminazene aceturate (44.44% w/w active ingredient) for the highest dosage of 40mg/kg bw was prepared by dissolving 90mg of the drug powder in 10mls distilled water to give a concentration of 4mg/ml, whereas Melarsoprol (5 ml vials of 180 mg) was first prepared by mixing (vortex) 1 Ml of the stock solution to 4 Ml of 50% propylene glycol to give a concentration of 7.2mg/ml (72mg/kg).This was further diluted to 40mg/kg by mixing(vortex) 5.6 ml of the 7.2mg/ml with 4.4mls of 50% propylene glycol to give a concentration of 4mg/ml (40mg/kg) The drug solutions for the 40mg/kg dose of each drug were then diluted serially using distilled water to give dosages for the 20, 10, 5, 2.5, 2, 1mg/kg.

### Statistical analysis

Analysis was carried out to test if there exists significant differences between the four classes of virulence using PP, parasitaemia progression, PCV, body weights changes and survival as the response variables. The data obtained from the study were summarized using descriptive statistics. General linear model in SAS was used to test significance at p<0.05 level, of differences between means of the 4 virulence classes. Survival data analysis was carried out employing the Kaplan–Meier method on StatView (SAS Institute, Version 5.0.1) statistical package for determination of survival distribution function. Rank tests of homogeneity were used to determine the effect of virulence on host survival time.[26].

## Results

### Survival time and classification

The 12 *T. b. rhodesiense* clones exhibited variations in survival time (Fig 1) and were classified into four classes of virulence based on these survival time data as shown (Table 1). A total of 3/12 clones were very acute, 3/12 were acute, 2/12 sub-acute and 4/12 chronic (Table 1). As expected, the shortest mean± SE survival time of 8.7 ± 0.2 days was observed in mice infected with the very-acute clones (Table 1). When the survival times of mice infected with isolates in the different virulence classes were compared, the Wilcoxon and Logrank tests p-value was 0.001. All control mice survived to the end of the experimental period of 60 days and their survival time data were therefore categorized as censored.

**Fig 1.**
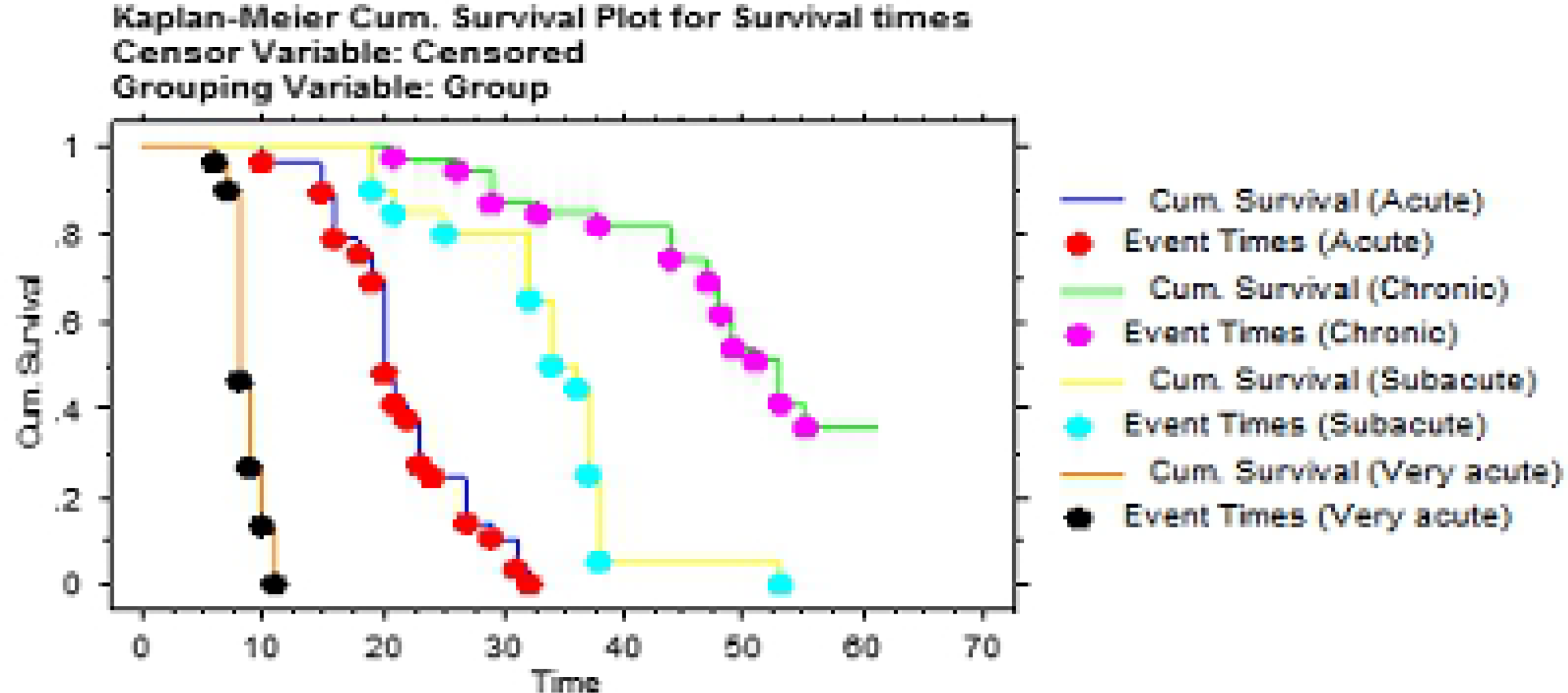
The survival times for mice infected with twelve *T. b. rhodesiense* clones. The clones were classified as very-acute (0-15 dpi), acute (16-30 dpi), sub-acute (31-45 dpi) and chronic *Tbr* (46-60 dpi); *dpi=days post infection*.

### Pre-patent period and parasitaemia progression

The mean ±SE pre-patent period in infected mice are summarized in (Table 1) as: very-acute *Tbr* clones: 4.7±0.09 dpi, acute *Tbr* clones: 5.0±0.2 dpi, sub-acute *Tbr* clones: 5.2±0.08 dpi and chronic *Tbr* clones: 6.3±0.2 dpi. Despite the apparent increasing trend of these data, these differences were however not statistically significant (p> 0.05). Summary analysis on mean peak parasitaemia (Mean ± SE) and number of days to peak parasitaemia (DPP) in each class are presented in (Table 1). Parasitaemia increased significantly (p<0.001) with days post infection for all the groups. However, when all virulence classes are compared, parasitaemia (Fig 2) was significantly (p<0.001) higher in mice infected with very-acute clones. In mice infected with very-acute clones, parasitaemia was characterized by a single wave (Fig 2) whereas parasitaemia progression in the other classes was characterized by two waves.

**Fig 2.**
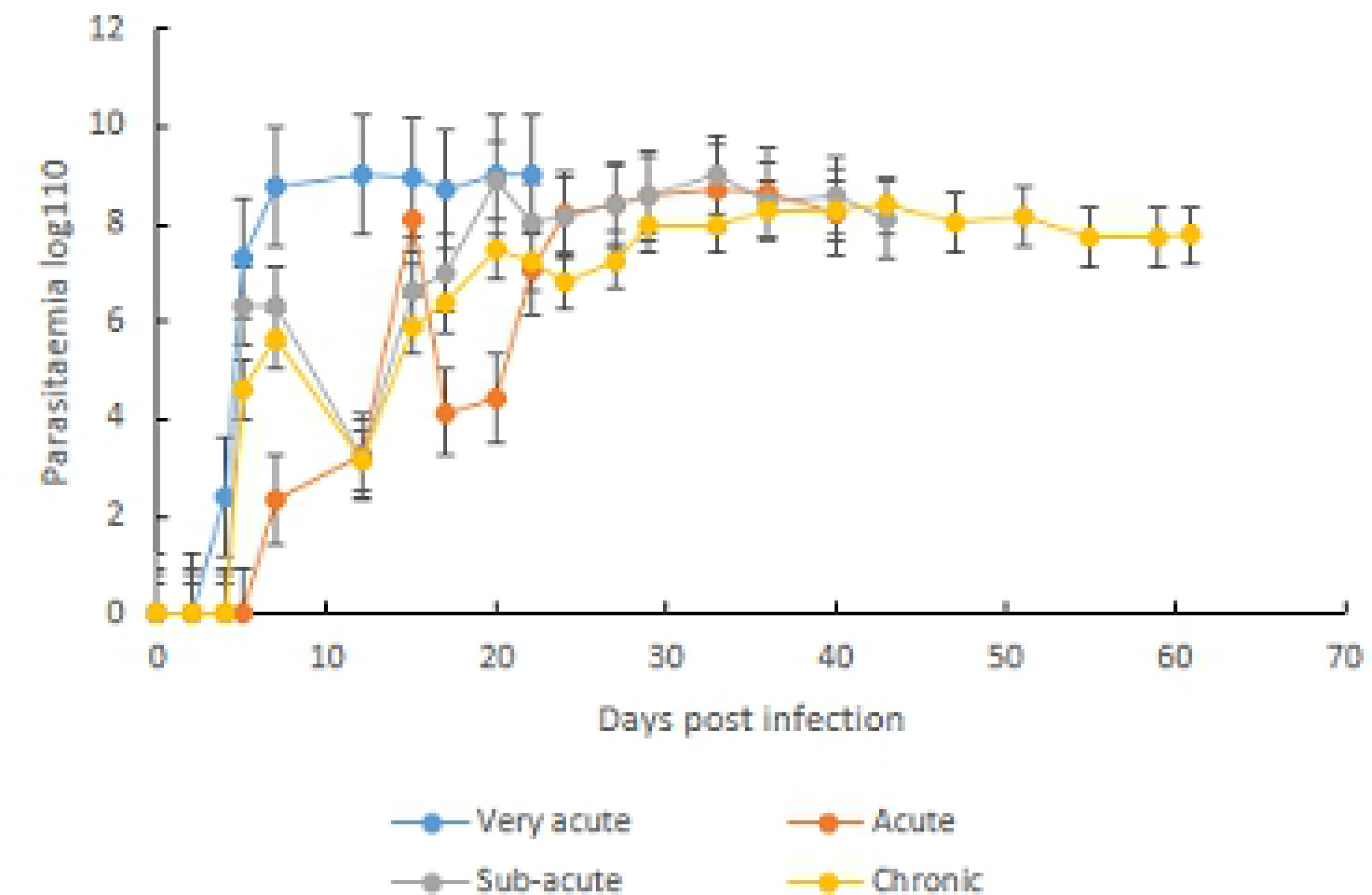
Parasitaemia progression in mice infected with the four classes of *T. b. rhodesiense* clones

**Table 1.** Changes in virulence biomarkers in mice infected with twelve *Trypanosoma brucei rhodesiense* clones.

| Class | Clone ID | Locality of isolation | Iso.<br>yr | PP | Peak<br>Para. | DPP | MST |
| --- | --- | --- | --- | --- | --- | --- | --- |
| very-acute | KETRI 2482 | Lumino, Uganda | 1969 | 5±0 | 1x10<sup>6</sup> | 8±0.2 | 9±0.4 |
|  | KETRI 3304 | Lugala, Uganda | 1971 | 5±0 | 7.6x10<sup>8</sup> | 7.8±0.4 | 9±0.4 |
|  | KETRI 3803 | Busia, Kenya | 1961 | 4±0 | 9.3x10<sup>8</sup> | 6.4±0.2 | 8.2±0.3 |
|  | **Group mean ±SE** |  |  | **4.7±0.9** | **8.9x10<sup>8</sup>** | **7.4±0.5** | **8.8±0.2** |
| Acute | KETRI 2487 | Busoga, Uganda | 1972 | 4±0 | 6.2x10<sup>8</sup> | 7±0 | 18.1±0.7 |
|  | KETRI 3800 | Busia, Kenya | 2000 | 5.2±0.1 | 1.2x10<sup>8</sup> | 6±0 | 26.0±2.0 |
|  | KETRI 3801 | Busia, Kenya | 1989 | 6±0 | 6.8x10<sup>7</sup> | 7.3±0.3 | 20.4±0.8 |
|  | **Group mean ±SE** |  |  | **5.03±0.16** | **1.7x10<sup>8</sup>** | **6.7±1.3** | **21.6±1.0** |
| Sub-acute | KETRI 3798 | Busia, Kenya | 1989 | 5.3±0.16 | 1.6x10<sup>8</sup> | 9.3±1.7 | 28.2±2.0 |
|  | KETRI 3926 | Busoga, Uganda | 1972 | 5±0 | 1.7x10<sup>8</sup> | 6.6±0.2 | 38.9±1.6 |
|  | **Group mean ±SE** |  |  | **5.2±0.8** | **1.5x10<sup>8</sup>** | **7.9±0.9** | **33.6±1.8** |
| Chronic | KETRI 3928 | Tororo, Uganda | 1992 | 6.3±0.3 | 7.9x10<sup>7</sup> | 7.0±0 | 51.8±4.1 |
|  | KETRI 3664 | Busia, Kenya | 1997 | 6.0±0 | 4.8x10<sup>7</sup> | 7.2±0.3 | 45.6±1.6 |
|  | KETRI 3380 | Busoga, Uganda | 2000 | 5.9±0.5 | 1.3x10<sup>8</sup> | 7.3±0.2 | 55.5±2.3 |
|  | KETRI 3305 | Lugala, Uganda | 1971 | 6.7±0.2 | 3.2x10<sup>8</sup> | 8.7±0.3 | 46.7±5.1 |
|  | **Group mean ±SE** |  |  | **6.3±0.2** | **1.1x10<sup>8</sup>** | **7.5±0.2** | **49.9±1.8** |

Key: PP-pre-patent period, Iso Yr-year of isolation, Par-parasitaemia, DPP -days to peak parasitaemia, MST-mean survival times

### Packed Cell Volume (PCV)

The pre-infection PCV data of for all infected and control mice groups (Fig 3) were not statistically different (p > 0.05). The PCV of the infected mice groups declined significantly (p < 0.001) with days post infection when compared with the PCV of the non-infected control mice which remained largely constant throughout the duration of the study (Fig 3). However, the onset and severity of the anemia, as shown by the decline in PCV, was most prominent for mice infected with the isolates classified as very acute (Fig 3). In these mice, the PCV declined significantly (p<0.001), from 49.7±0.8 at baseline (day 0) to 26.0±0.5 at 14 dpi equivalent to 47.7% decline. The lowest infection-related decline in PCV (Fig 3) was recorded in the mice infected with isolates classified as chronic clones, with the PCV declining from 49.6±0.9at baseline to 43.5±1.0at 14 dpi (12.3%).

**Fig 3.**
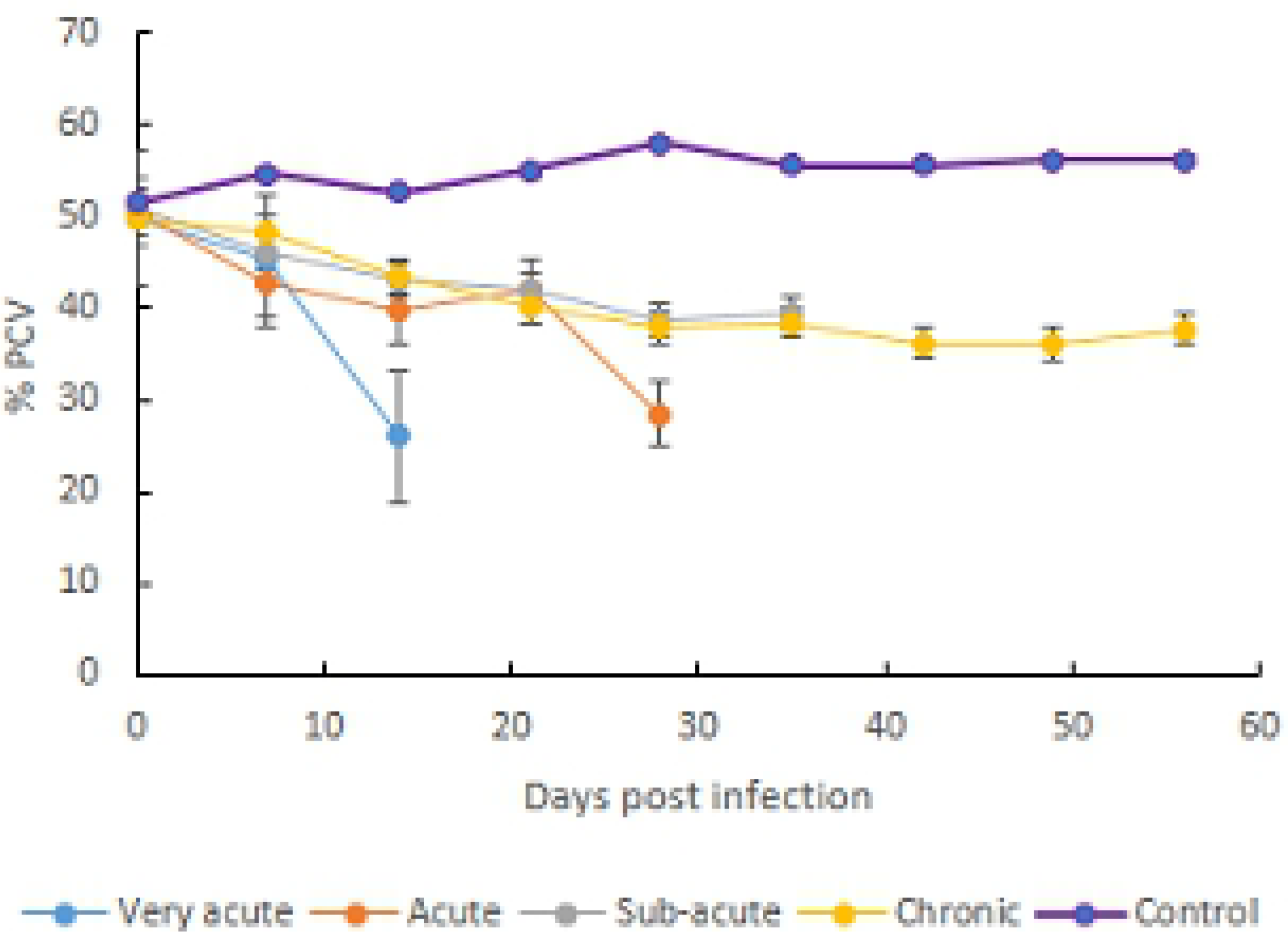
Mean ± SE PCV decline in mice infected with *T.b. rhodesiense* very-acute isolates, acute isolates, sub-acute isolates and chronic isolates clones.

### Body weight

The mean ±SE pre-infection body weight data for infected and control groups (Fig 4) were not statistically different (p > 0.05). Between 7 and 14 days post infection, all infected mice groups exhibited a decline in mean body weight (Fig 4) while the body weight of un-infected control mice did not change (Fig 4). However, mice groups infected with isolates classified as acute, subacute and chronic exhibited recovery of their body weights starting 14 dpi. Mice group infected with isolates in the very acute virulence class did not survive beyond 14 dpi (Fig 4).

**Fig 4.**
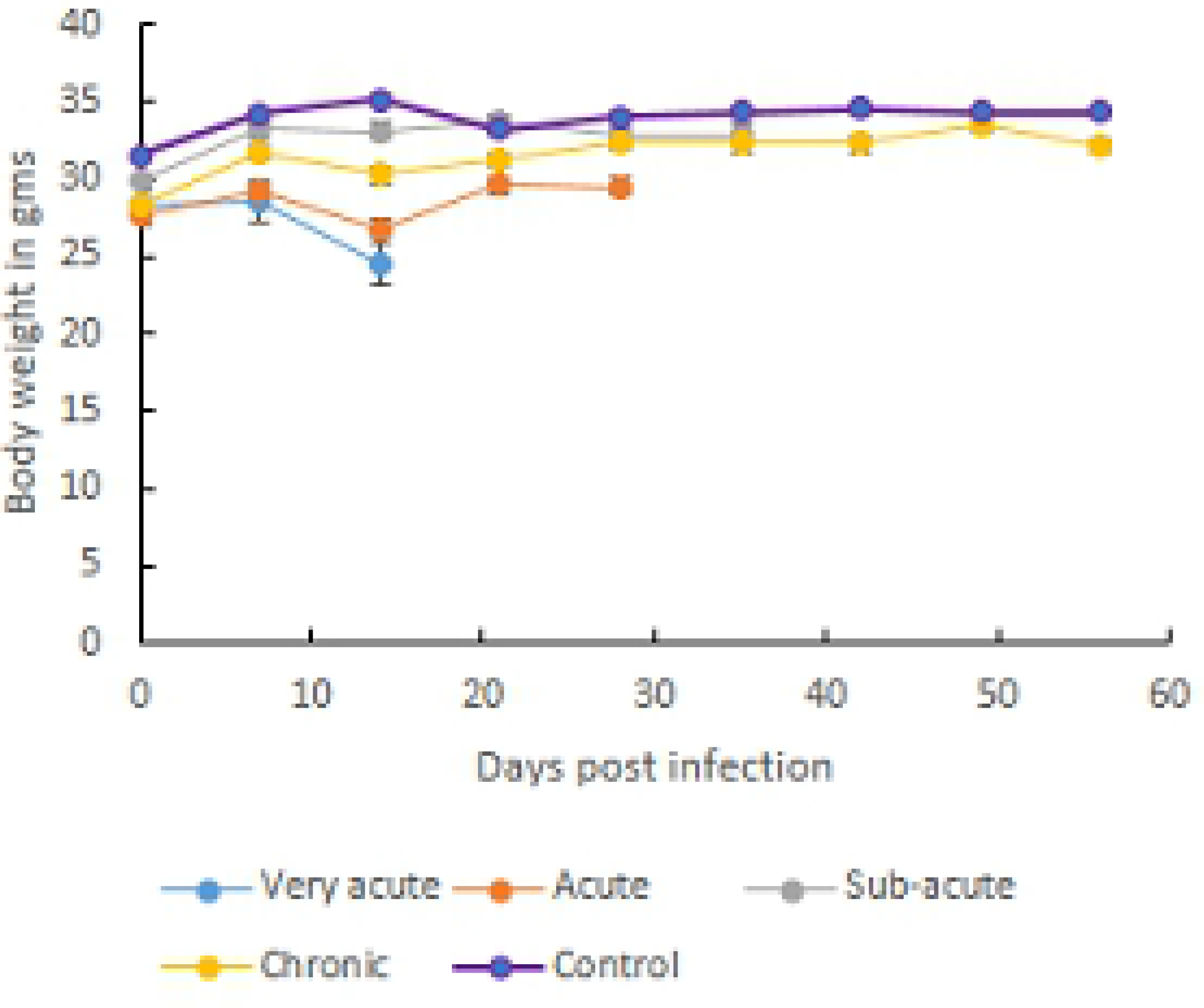
Mean ± SE body weight changes in mice infected with *T.b. rhodesiense* very-acute isolates, acute isolates, sub-acute isolates and chronic isolates clones.

### Drug sensitivity results

The results of the drug-sensitivity testing for the reference sensitive (*Tbr* KETRI 3738) and drugresistant (Tbr KETRI 2538) isolates are shown in (Table 2). The reference drug-resistant isolate was confirmed to be resistant to melarsoprol, Pentamidine and Diminazene aceturate at dose rates ranging from 1-20 mg/kg body weight (Table 2). However, infected mice were cured with all three drugs at a dose rate of 40 mg/kg body weight. With respect to suramin, the reference resistant isolate was sensitive to all doses equal to or greater than 5 mg/kg body and is therefore characterized as sensitive (Table 2). On the other hand, reference drug-sensitive isolate was confirmed to be sensitive to all doses of melarsoprol, ranging from 1-40 mg/kg bwt. It was also fully sensitive to diminazene aceturate at dose rates ranging from 2.5-40mg/kg bwt. The reference sensitive isolate was sensitive to pentamidne at all doses above 4 mg/kg bwt (Table 2). It was also sensitive to all doses of suramin equal to or greater than 2.5 mg/kg (Table 2)

The results of drug sensitivity experiments for the test *Tbr* clones are summarized in (Table 3). All the isolates recorded at least 80% cure rates to all the drug dose regimens evaluated in this study (Table 3) and were therefore classified as sensitive. However, a few cases of relapses were observed in 1/5 (20%) mice infected with KETRI 2482 (very-acute group) and treated with diminazene aceturate at 2.5mg/kg, and KETRI 2487 (acute) and KETRI 3926 (sub-acute) treated with pentamidine at 5mg/kg. In mice infected with KETRI 3928 (Chronic), 4/5 treated with diminazene aceturate at 2.5mg/kg and 5/5 treated at 20 mg/kg died at 47 days post treatment due to causes not related to trypanosome infection (Table 3). No relapses were observed in mice groups that were treated with either melarsoprol (1 and 20 mg/kg) or suramin at 2.5 mg/kg (Table 3)

**Table 2.** Results of drug sensitivity evaluation of reference KALRO-BioRI sensitive and resistant *T b rhodesiense* isolates.

| *Sensitive Isolate KETRI 2537* |  |  |  | *Resistant isolate KETRI 2538* |  |  |
| --- | --- | --- | --- | --- | --- | --- |
| Drug | Drug dose (mg/Kg) | Mice cured/5 | Status | Drug dose (mg/Kg) | Mice cured/5 | Status |
| MeIB | 40 | 5 | (s) | 40 | 5 | **(S)** |
|  | 20 | 4 | (s) | 20 | 0 | (R) |
|  | 10 | 5 | (s) | 10 | 0 | (R) |
|  | 5 | 5 | (s) | 5 | 0 | (R) |
|  | 2.5 | 5 | (s) | 2.5 | 0 | (R) |
|  | 1 | 5 | (R) | 1 | 0 | (R) |
| diminazene aceturate | 40 | 5 | (s) | 40 | 5 | **(S)** |
|  | 20 | 5 | (s) | 20 | 0 | (R) |
|  | 10 | 4 | (s) | 10 | 0 | (R) |
|  | 5 | 5 | (s) | 5 | 0 | (R) |
|  | 2.5 | 5 | (s) | 2.5 | 0 | (R) |
|  | 1 | 2 | **(R)** | 1 | 0 | (R) |
| Pentamidine | 40 | 5 | (s) | 40 | 5 | **(S)** |
|  | 20 | 5 | (s) | 20 | 1 | (R) |
|  | 10 | 5 | (s) | 10 | 1 | (R) |
|  | 5 | 4 | (s) | 5 | 0 | (R) |
|  | 2.5 | 2 | **(R)** | 2.5 | 0 | (R) |
|  | 1 | 0 | **(R)** | 1 | 0 | (R) |
| Suramin | 40 | 5 | (S) | 40 | 5 | **(S)** |
|  | 20 | 5 | (S) | 20 | 5 | **(S)** |
|  | 10 | 5 | (s) | 10 | 5 | **(S)** |
|  | 5 | 5 | (s) | 5 | 5 | **(S)** |
|  | 2.5 | 4 | (s) | 2.5 | 2 | (R) |
|  | 1 | 1 | **(R)** | 1 | 0 | (R) |
| Control | - | 10 |  | - | 10 |  |

Key:- Not treated; The mice groups (n=5) were treated 24hours post inoculation with the isolates and monitored for 60 days post treatment. An isolate is coded as sensitive (S) when at least 4/5 mice survived for at least 60 days without trypanosome relapse. All other results are coded as resistant (R).

**Table 3.**
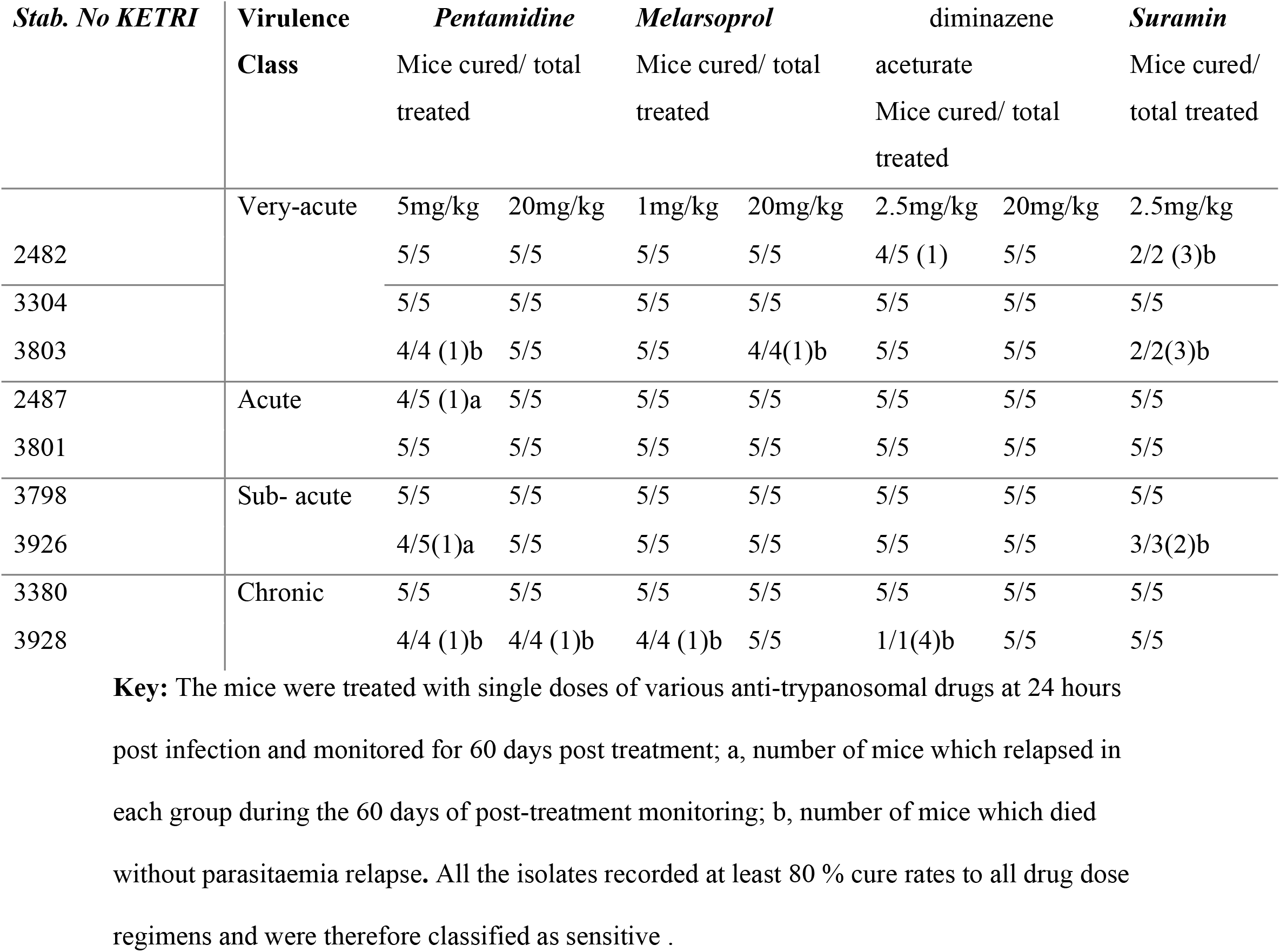
Results of drug sensitivity evaluation of *T b rhodesiense* clones in the mouse model.

## Discussion

In this study, we characterized the virulence and anti-trypanosomal drug sensitivity patterns of 12 *T. b. rhodesiense* cloned stabilates. We used *T. b rhodesiense* clones because they represent a homogeneous population of genetically identical trypanosomes [27]. The results demonstrated the existence of variations in virulence of *T. b. rhodesiense* cloned stabilates (Table 1) which is interesting because all study isolates were originally recovered from western Kenya and eastern Ugandan, regions that are considered to belong to the same Busoga focus of HAT. While it is a well established fact that clinical profiles of HAT patients in eastern Africa Uganda and Kenya differ from those of patients in Southern African HAT foci such as Malawi and Zambia [8] our study suggests these differences would be expected to be present even within a single HAT focus. Our results are in agreement with a study by [28] on a number of isolates from eastern Uganda in mice which showed that distinct acute and chronic strains of *T. b. rhodesiense* circulated in the focus. They are also in agreement with previous reports for *T. b. gambiense* isolates [29].

We used mean survival time (MST) of mice post-infection as the main indicator of virulence as previously reported [23,30]; [24] and observed that the *Tbr* isolates were well distributed among the four virulence classes. This finding explains why clinical syndromes in HAT patients differ significantly even in a single HAT focus, thus complicating HAT diagnosis. Infective isolates that allowed mice to have long survival times, hence chronic infections, may indicate presence of enriched population of stumpy forms which aids in prolonging host survival and enhancing the probability of parasite transmission [31]. The mean survival times for the very acute clones was 8.7 days suggesting the hosts were overwhelmed by the first parasitaemia peak before the proliferating slender forms differentiated into short stumpy forms [32]. The majority of the *Tbr* clones used in this study had undergone a minimum number of passages since isolation (Table 2) confirming therefore that the observed differences in isolate virulence is an intrinsic attribute as previously reported [33].

Parasitaemia progression among *Tbr* isolates assigned to different virulence classes on the basis of survival time were significantly different. This finding is in agreement with previous studies in which virulence of different species of trypanosomes was characterised using parasitaemia, intensity of anaemia (PCV) and weight loss experienced by the host during the infection period [24]. In our study, parasitaemia of isolates in the very-acute virulence class was represented by a single wave whereas the acute, sub-acute and chronic virulence classes were represented by two waves. (Fig 2). This is in agreement with studies by [34] who observed that acute infections results from uncontrolled proliferation of the slender trypanosome forms without differentiation into short stumpy forms and hence kills the host before tsetse transmission takes place [34]. In contrast, chronic infection is characterized by appearance of progressive waves of parasitaemia, with each distinct wave being composed of trypanosomes with antigenically distinct coats, and with parasites easily differentiating into the transmissible short stumpy forms. This perhaps explains why highly virulent trypanosomes are not easily transmissible as was observed by[19] that tsetse flies infected with chronic *T. b. brucei* recorded highest mature infection as opposed to those infected with highly virulent trypanosomes. Our results are important as they reveal that majority of *T. b. rhodesiense* infections are in the bracket of (acute, sub-acute and chronic) classes of virulence and can easily be transmittable.

In the present study, all infected mice recorded a decline in PCV signifying the development of *T. b. rhodesiense* induced anemia. Our observation was in agreement with previous studies which reported anaemia as a key feature both in humans [35] and in the monkey model [6]. As with parasitaemia and survival time parameters, the development of anemia was significantly pronounced in mice infected with very-acute clones. This finding is consistent with observations by [36] who reported that acute infection of mice with *Trypanosoma cruzi* was characterized by an exponential growth of parasites and high mortality accompanied by anemia. A similar observation was made by [24] in mice infected with *Trypanosoma evansi*. In contrast, anaemia in mice infected with clones in the other various classes of virulence (acute, sub-acute and chronic) stabilized or recovered characteristic of the chronic phase anaemia [37]. The severity of anemia is determined by parasite virulence, time lag from infection to therapeutic intervention and individual host differences [38].

Our results showed on body weight showed a decline in the early days of infection (7-14 dpi) which thereafter recorded recovery with exception of very-acute infected mice. This decline was however not significant. Our observation is important as it confirms previous observation [19] that body weight alone cannot conclusively serve as a virulence biomarker. Previous authors [39] attributed decline in body weight to reduced food intake. In our study, we did not measure the food intake. The failure by infected mice to register a decline calls for further investigation on causes of body weight changes in trypanosomes infected animals and especially after previous studies have recorded an increase in body weight in *T.evansi* [24] and in *T. b. brucei* or *T. congolense* [19] infected mice with days post infection.

Our results on drug sensitivity tests showed that all the study isolates were sensitive to melarsoprol, pentamidine, diminazene aceturate and suramin. The sensitivity of these isolates to suramin and melarsoprol is significant since these are the drugs which are recommended by WHO (2018) to treat early and late stages of *Tbr* HAT respectively. On the other hand the sensitivity of the *Tbr* isolates to diminazene aceturate, is an indicator of the utility of these drug when administered to livestock reservoirs of *Tbr* isolates as practiced in disease HAT control programmes in endemic countries [40] Interestingly, however, the single cases of relapses encountered in mice infected with KETRI 2482 (very-acute virulence class), KETRI 2487 (acute virulence class) and 3926 (sub-acute virulence class) were all against the two diamidines (pentamidine or dimainazene) but not against suramin or melarsoprol (Table 3) which is consistent with clinical practice of not using these specific diamidines to treat *Tbr* HAT (WHO, 2018). Overall, the fact that the test isolates were all sensitive (at least 80% cure rates) to the drugs suggests there was no relationship between isolates’ virulence and their sensitivity to antitrypanosomal drugs.

The KALRO-BioRI reference isolate considered to be drug resistant was confirmed in this study to be resistant to melarsoprol, pentamidine and diminazene aceturate (Table 2). In general drug resistance is attributed to reduced drug uptake due the mutation or absence of drug uptake gene [41] as well as by enhanced drug export, mediated by a multidrug resistance-associated protein,[42]. The uptake of the three drugs, melarsoprol, pentamidine and diminazene is mediated by the P2 transporter [12,43,44] which explains why resistance to all three drugs is linked. In contrast, uptake of suramin by trypanosomes is not mediated by the P2 transporter, hence the reason why the trypanosome, KETRI 2538, retains sensitivity to suramin

In summary, this study has found that there exists variations in virulence of isolates recovered from western Kenya/eastern Uganda HAT focus. Virulence is attributed to the production by the blood stream forms of membranous nanotubes that originate from the flagellar membrane and disassociate into free extracellular vehicles (EVs). This (EVs) contain several flagellar proteins that contribute to virulence [45]. Our results are important as they have demonstrated that virulence is not a hindrance in the control of trypanosomiasis by chemotherapy. However, our study only tested the drug sensitivity at 24 hours post infection before trypanosomes could establish themselves. There will be need to confirm our observation by administering the drugs when animals are parasitaemic to ascertain the effectiveness of the drug in clearing the established infections.

## Acknowledgment

We acknowledge the Director, KALRO for permission to publish this study. Our other acknowledgment goes to Dr. Johnson Ouma, former Center Director (Trypanosomiasis Research Center) BioRI for supervision and facilitation, technical staff of KALRO-BioRI and in particular John Ndichu, Jane Hanya for taking care of the infected mice. Gilbert Ouma and Mr. Mageto for the preparation of drugs

